# CINNA: An R/CRAN package to decipher Central Informative Nodes in Network Analysis

**DOI:** 10.1101/168757

**Authors:** Minoo Ashtiani, Mehdi Mirzaie, Mohieddin Jafari

## Abstract

In network science, usually there is a critical step known as centrality analysis. This is an important step, since by using centrality measures, a large number of vertices with low priority are set aside and only a few ones remain to be used for further inferential outcomes. In the other words, these measures help us to sieve our large network and distinguish coarse vertices. By that, important decisions could be made based on the circumstances of these vertices on the overall behavior of networks. These vertices are potentially assumed as central or essential nodes. However, the centrality analysis has always been accompanied by a series of ambiguities, since there are a large number of well-known centrality measures, with different algorithms pointing to these essential nodes and there is no well-defined preference. Which measure explore more information in a given network about node essentiality according to the topological features? While here, we tried to provide a pipeline to have a comparison among all proper centrality measures regarding the network structure and choose the most informative one according to dimensional reduction methods. Central Informative Nodes in Network Analysis (CINNA) package is prepared to gather all required function for centrality analysis in the weighted/unweighted and directed/undirected networks.

**Availability and implementation:** *CINNA* is available in CRAN, including a tutorial. URL: https://cran.r-proiect.org/web/packages/CINNA/index.html

## 1 Introduction

Centrality measure is a mathematical definition of a node position through a network. By the development of network science, several centrality measures are proposed regarding to the various notions, e.g. distance, degree, eigen and neighborhood. However, the critical concern is, which centrality measure can precisely point at the influencer nodes of a network. At 2006, Dwyer et al. used visualization methods like orbit and hierarchy based comparison to assess the difference among centrality measures (Dwyer, Hong et al. 2006). At the same time, Borgatti and Everett realized that a network does not subtend separate cohesive subgroups, so measures of centrality are tended to be closely connected with the cohesive subgroup of a network structure (Borgatti and Everett 2006). Thereafter, some researchers tried to compare centralities by means of the correlation between them (Valente, Coronges et al. 2008, Li, Li et al. 2015).

While choosing an appropriate centrality is riddling with controversy for many years, in a companion work, we introduced a new approach to pick the most informative one within a network according to dimensional reduction methods. In brief, we analyzed 27 centrality measures in 14 distinct networks to show that properly detecting central nodes requires utilizing methods such as principal components analysis (PCA) (Ashtiani, et al., 2017). To best of our knowledge, no specified pipeline has been introduced for prioritizing informative centrality measures regarding the network topology. Here, we introduce CINNA R package, which is a collection of functions to calculate, compare, assort, and visualize network centrality measures in a comprehensive manner.

## 2 Methods

### 2.1 Input: network component extraction

The first step, before the centrality analysis, is extracting the giant component of a given network as most of the centrality measures are calculable on the connected component of a network. Hence, functions for segregating components of a network are available in various graph formats such as igraph (Csardi and Nepusz 2006), network (Butts 2008), adjacency matrix or edge list. In addition to all those formats in a single command, *CINNA* has also provided extra functions for bipartite graphs, which able users to apply centrality analysis on each projection that they want.

### 2.2 Calculation of centrality measures

In the next step, functions for indicating appropriate centrality types based on the network structure i.e. undirected-unweighted, undirected-weighted, directed-unweighted and directed-weighted graph, are provided. By finding out the calculable proper centrality, user can specify what centrality types he wants to compute and have a comparison between them. Then, desired measures are computed in a particular function.

### 2.3 Determination of most informative centrality measure

Thirdly, the fundamental analysis i.e. determination of most informative centrality measure can be applied. *CINNA* has prepared two diverse procedure as Principal Component Analysis(PCA) (Abdi and Williams 2010) and t-Distributed Stochastic Neighbor Embedding (t-sne) algorithm (Maaten and Hinton 2008) to distinguish most informative centralities. Both of them are dimensionality reduction approaches regarding linear and non-linear analysis. In PCA method, with respect to the contribution criteria, highly informative centralities concerning principal components can be established and visualized. By identifying the most informative one, we can infer which centrality type best fits our network and can notices essential nodes more accurately (Ashtiani, Salehzadeh et al. 2017). In the nonlinear procedure, t-sne algorithm, the cost criteria is exploited to acquire the most informative one. Hence, a user can be able to compare computed centrality results based on what algorithm he/she assigns.

### 2.4 Visualization of centrality analysis

In addition to supplying a suitable pipeline for designating the most informative centrality, *CINNA* package gathers diverse visualization techniques for comparing results of centrality analysis including heatmap, dendrogram and pairwise scatter plot. These methods are prepared to facilitate the analysis and quantification of the centrality results and the overlap among the diverse groups. Moreover, the whole network can be illustrated by fixing a centrality type whereas the shape sizes of vertices indicate the centrality values.

## 3 Example of use

We performed the approach on several datasets and confirmed that the functionality of the procedure improves accuracy of the centrality analysis. The unified manual can be accessed using the R command help (package = CINNA). Besides, an immense step-by-step tutorial on all substantial features of the package can be approached using the command browseVignette (“CINNA”). An example usage in R is shown in the following, where we employed Zachary network (Zachary 1977) to glance on CINNA functionality in a nutshell.

~~~
# Install the package
install.packages("CINNA")
# Load the package
library(CINNA)
# Load the data set
data(“zachary”)
# Indicate the names of proper centralities regarding the newtork structure
properCentralities <-proper_centralities(zachary)
# Calculation centrality measures
centralitiesValues <-calculate_centralities(zachary, include = c(“Closeness Centrality
(Freeman)”, “Decay Centrality”, “Degree Centrality”, “Radiality Centrality”))
# Apply PCA on the computed centrality values and visualize most informative ones
pca_centralities(centralitiesValues)
~~~

The corresponding results with additive visualization plots are illustrated in Figure 1. As shown, Decay centrality (Toropov 2017) have the highest level of contribution value among our small collection.

**Fig. 1.**
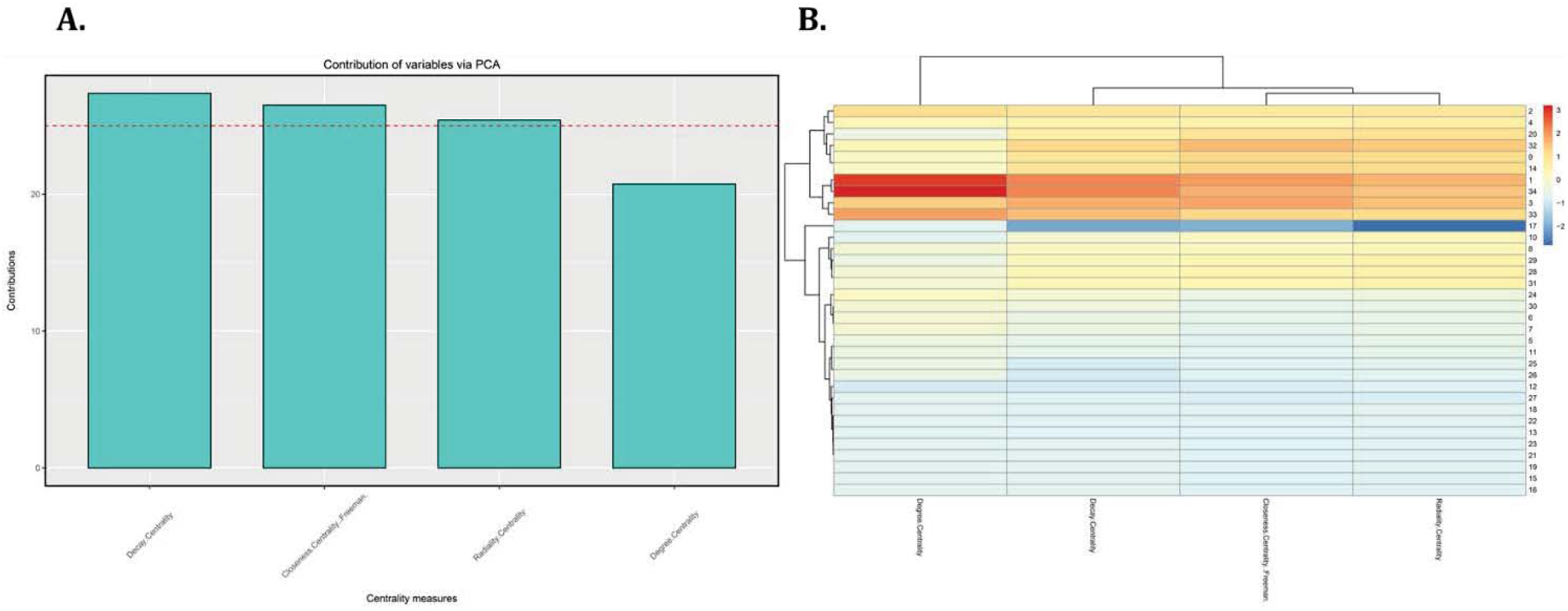
Exemplary uses of CINNA on Zachary network. A) PCA contribution bar plot. The decay centrality is the most informative one unlike the degree centrality. B) Heat map of nodes centrality measure values. Colors spectra from blue to red represent nodes that have lowest to highest centrality values.

## 4 Conclusions

The CINNA package currently includes 23 user-level functions along with five real network examples (Zachary 1977, Felleman and Van 1991, Barneh, Jafari et al. 2016, Dataset 2016) which helps user to have a different experience from the centrality network analysis. An essential part of the advancement process is the researcher’s feedbacks. We appreciate receiving all suggestions and comments from the users for the next version.

## Acknowledgements

The authors appreciate Reza Karbalaei for designing the logo of CINNA package, which can be seen http://iafarilab.com/cinna-page.html. Also, we acknowledge Ali Salehzadeh-Yazdi and Mahdi Jalili for suggestions and feedback.

